# Derivation of primed sheep embryonic stem cells and conversion to an intermediate naïve-like state

**DOI:** 10.1101/2025.11.22.689923

**Authors:** TS Shyamkumar, Manuel A. Vasquez-Hidalgo, Viju V. Pillai

**Affiliations:** Department of Comparative Pathobiology, College of Veterinary Medicine, Purdue University, West Lafayette, IN 47907, USA; Department of Veterinary & Biomedical Sciences, College of Agriculture, Food, and Environmental Sciences, South Dakota State University, Brookings, SD 57007, USA; Wille M. Reed Animal Disease Diagnostic Laboratory, Purdue University, West Lafayette, IN 47907, USA

**Keywords:** Sheep, embryonic stem cells, pluripotency, reporter

## Abstract

Embryonic stem cells (ESCs) derived from the inner cell mass of embryos possess unlimited self-renewal and pluripotency, offering a powerful system to study early development and enable genetic and biotechnological innovation. Although several livestock ESC lines have been reported in recent years, defining culture conditions that support stable long-term self-renewal and controlled transitions across pluripotent states remains challenging. Here, we report the *de novo* derivation of sheep embryonic stem cells (sESCs) from in vivo blastocysts using a chemically defined culture system. The derived cells exhibit morphological and molecular features of primed pluripotency and can be propagated under both feeder-dependent and feeder-free conditions without loss of identity or karyotypic stability. Building on this foundation, we developed enhancer-driven reporter lines that faithfully reflect OCT4 and SOX2 transcriptional activity, enabling dynamic visualization of pluripotency and differentiation in live cultures. These reporter systems revealed the responsiveness of sESCs to signaling modulation and provided a functional readout of pluripotency state transitions. When cultured in defined media previously shown to stabilize naïve pluripotency in human ESCs, sESCs adopted dome-shaped colony morphology, maintained OCT4, SOX2, and NANOG expression, retained differentiation potential, and exhibited a transcriptomic profile consistent with resetting to an intermediate pluripotent state with naïve-like morphological features. These findings establish stable sheep ESC lines and demonstrate their plasticity across the pluripotency spectrum, providing a valuable platform for investigating ruminant stem cell biology and advancing livestock biotechnology.

## Introduction

Embryonic stem cells (ESCs) derived from the inner cell mass of blastocyst-stage embryos possess unlimited self-renewal capacity and the ability to differentiate into all three germ layers [1]. These defining properties make ESCs invaluable for developmental biology, regenerative medicine, and biotechnology. In livestock, ESCs could be beneficial to animal agriculture by enabling precise genome editing, accelerating genetic improvement, and serving as renewable cell sources for cellular agriculture, including cultured meat and high-value proteins [2,3]. When combined with CRISPR-Cas9 technology and somatic cell nuclear transfer, livestock ESCs could provide a platform for generating animals with desirable traits and enhanced disease resistance [4]. Beyond agriculture, ESCs also serve as models for studying cellular differentiation, drug discovery, and toxicology, extending their utility into biomedical contexts [5].

Significant progress has been made in establishing pluripotent stem cells (PSCs) in rodents and primates, where distinct states of pluripotency, including naïve, primed, and intermediate including region specific, or formative states, have been captured and extensively characterized [6–11]. Naïve PSCs correspond to the pre-implantation epiblast and possess robust developmental potential, while primed PSCs are shown to have more restricted chimeric capacity. More recently, intermediate pluripotent states bridging naïve and primed have been identified in both mouse and human, highlighting the continuum of pluripotency [7,8,11]. Importantly, human and mouse models have also benefited from the development of advanced tools, such as enhancer-driven reporters, enabling real-time monitoring of pluripotency and facilitating detailed dissection of the molecular mechanisms underlying self-renewal and differentiation [12,13].

In livestock, however, progress has been slower, hindered by species-specific requirements [14]. Expanded potential stem cells (EPSCs) and primed ESCs have been described in cattle and pigs [15–18], yet reproducible derivation and long-term maintenance of other pluripotent states remain elusive. In sheep, only primed pluripotent stem cells have been convincingly established, with no reports of naïve or intermediate states resembling those in rodents or primates [19,20]. Furthermore, livestock ESC systems lack robust molecular tools for monitoring pluripotency, limiting precise manipulation of stem cell states. These challenges underscore the need to refine culture conditions, dissect signaling pathways, and establish molecular markers that define pluripotency in ruminants.

In this study, we sought to address these challenges by deriving and characterizing sESCs from in vivo blastocysts under chemically defined conditions. We evaluated their molecular and functional properties, including self-renewal, differentiation potential in vitro and in vivo, and karyotypic stability. We further generated enhancer-driven reporter lines to monitor pluripotency and examined their signaling pathway dependence. Finally, we tested conditions for reverting sESCs and established an intermediate state pluripotent stem cells that exhibit morphological, molecular, and transcriptomic features closer to the naïve state. Our findings provide new insights into sheep pluripotency and establish a foundation for advancing livestock stem cell biology.

## Materials and methods

All animal procedures were approved by the Institutional Animal Care and Use Committee, protocol number 0824002522 and conducted in accordance with guidelines.

### Culture of primed sheep embryonic stem cells

For the derivation of sESCs, embryos were plated on mitomycin-C-inactivated mouse embryonic fibroblasts (iMEFs) in a chemically defined NBFR medium [21]. This medium was prepared by combining equal volumes of DMEM/F-12 and Neurobasal medium, supplemented with 1% BSA (MP Biomedicals), N2 supplement and B27 supplement, GlutaMAX™ (Gibco), penicillin-streptomycin, and non-essential amino acids. In addition, the medium was added with 20 ng/mL Activin A (Peprotech, 120-14P), 20 ng/mL bFGF (Peprotech, 100-18B), and 3 µM endo-IWR-1 (Biogems, 1128234). All components were mixed under sterile conditions, and the medium was stored at 4°C and used within one week. For feeder free culture, cells cultured on iMEFs were sequentially adapted. Medium was replaced daily, and passaging was performed every 2–3 days using mechanical dissociation of colonies with either a stem cell passaging tool (StemPro EZPassage™, Invitrogen) or a pipette tip. A Rho-associated kinase (ROCK) inhibitor, Y-27632 (10 µM), was added to the medium during passaging to enhance cell survival. Cultures were maintained at 37°C in a humidified incubator with 5% CO₂ under ambient oxygen conditions. The cells were cryopreserved in a freezing medium composed of 90% NBFR and 10% dimethyl sulfoxide.

Sheep embryonic stem cells were also cultured in a modified mTeSR-based medium referred to as emTeSR. This medium was prepared by combining mTeSR1 basal medium and supplement (StemCell Technologies), supplemented with GlutaMAX™, penicillin-streptomycin, and MEM non-essential amino acids and 20 ng/mL Activin A, 20 ng/mL bFGF, and 3 µM endo-IWR-1, 2.5 µM IWP1 or 2.5 µM XAV-939. All the components were mixed under sterile conditions, stored at 4°C and used within one week. For the feeder-free culture, Matrigel® (Corning), Geltrex (Gibco), Laminin-521 (Gibco), collagen IV (Corning) and vitronectin (Gibco) were used as per the manufacturer’s instructions.

### Conversion of sheep ESCs using different media conditions

Primed sESCs grown on NBFR or emTeSR were changed into different defined naïve pluripotency media. All formulations were based on a 1:1 mixture of DMEM/F-12 and Neurobasal medium, supplemented with 1% BSA, N2, B27, GlutaMAX™, penicillin-streptomycin, and non-essential amino acids with added supplements as given below. Cells were maintained on iMEFs or feeder-free substrates. HENSM [22] had 20 ng/mL human LIF, 20 ng/mL Activin A, 1 µM PD0325901, 2 µM XAV-939, 2 µM Go6983, 1.2 µM CGP77675, and 50 µg/mL ascorbic acid. 5i/L/A medium [23] was prepared using 20 ng/mL LIF, 10 ng/mL Activin A, 1 µM PD0325901, 1 µM IM-12, 0.5 µM SB590885, 1 µM WH-4-023, 10 µM Y-27632, and 0.1 mM 2-mercaptoethanol. The 2i/LIF medium [24], contained 0 ng/mL LIF, 1 µM PD0325901, and 1 µM CHIR99021, whereas the t2iL+Gö medium [25] contained 10 ng/mL LIF, 1 µM PD0325901, 1 µM CHIR99021, and 2 µM Go6983.

### Alkaline phosphatase staining and immunocytochemistry

Staining for ALP activity was performed using Vector Alkaline Phosphatase Substrate kit as per manufacturer instructions (Vector labs). All steps involved in immunolabeling and imaging have been previously described [26]. Briefly, cells were cultured on glass coverslips and fixed by adding 4% paraformaldehyde directly to the culture medium for 15 min, followed by replacement with fresh fixative for another 15 min. After PBS washes, cells were permeabilized with 0.5% Triton X-100 for 15 min and blocked with 0.5% goat serum for 1 h at room temperature. Primary antibodies (1:200 in goat serum) were applied for 3 h at room temperature or overnight at 4°C, followed by Alexa Fluor-conjugated secondary antibodies (1:1000 in PBS) for 1 h in the dark. Coverslips were mounted with ProLong™ Gold Antifade Mountant containing DAPI (Invitrogen, P36941), and images were acquired using either a Leica fluorescence microscope or a Zeiss confocal microscope. Antibodies used include Sox2 (Invitrogen); Oct4 (Invitrogen), Pancytokeratin (Zeta), Vimentin (Zeta), FOXA2 (Invitrogen) and H3K27me3 (Invitrogen).

### Teratoma assay

Cells were dissociated into single-cell suspensions with TrypLE and mixed on ice with cold Matrigel® diluted in DMEM/F12, yielding approximately 1 × 10^6^ cells in 200 µl. The suspension was transferred into a pre-chilled 1 ml syringe fitted with a 30G needle. Six- to eight-week-old immunodeficient NSG mice (NOD.Cg-Prkdc^scid^Il2rg^tm1Wjl^/SzJ, Jax^®^ mice; Jackson Laboratory) were injected subcutaneously at two to three sites along the flank or dorsal region, with 50–100 µl delivered per site. Animals were maintained under standard housing conditions and regularly monitored for teratoma development. At six weeks post-injection, mice were euthanized, and the resulting teratomas were harvested, fixed in 4% formaldehyde, and processed for paraffin embedding. 4 µm sections were cut and stained with hematoxylin and eosin using standard histological procedures reported previously [27].

### Embryoid body formation and in vitro trilineage differentiation

Embryoid body formation was based on previously reported protocols [28]. Briefly, undifferentiated primed embryonic stem cell colonies were mechanically dissected into medium-sized fragments with a fire-drawn glass Pasteur pipette and seeded into 24-well low-attachment plates. After five days, embryoid bodies were transferred to 0.1% gelatin-coated dishes and cultured in spontaneous differentiation medium [DMEM/F-12 with 20% FBS, 2 mM L-glutamine, 1% NEAA, and 0.1 mM 2-mercaptoethanol], with half the medium replaced every other day. After 14 days, cells were stained for lineage-specific markers.

### Karyotyping

For chromosome spread preparation, colcemid (KaryoMAX™, Gibco; 10 µg/mL) was added to the culture medium at 20 µL per 1.5 mL and incubated for 1.5 hours. Cells were rinsed with PBS, dissociated using TrypLE Express to obtain a single-cell suspension, and pelleted by centrifugation at 122 × g for 5 min. The pellet was resuspended in 5 mL pre-warmed 75 mM KCl hypotonic solution, incubated at 37°C for 15 min, and fixed with methanol:acetic acid (3:1) added in steps (3 mL + 2 mL). Fixed cells were stored overnight at 4°C. Drops of the suspension were applied onto clean, chilled microscope slides, air-dried for 10–15 min, and stained with Giemsa solution for 10 min. Slides were rinsed in deionized water, air-dried, mounted in mounting medium, and visualized under oil immersion at 100× magnification.

### RNA Sequencing

Two primed sESC lines cultured in NBFR media under feeder-free conditions were used for RNA sequencing (RNA seq). Similarly, naïve-like cells cultured feeder free conditions were used. RNA was extracted with the RNeasy Plus Micro Kit (Qiagen), and polyA-enriched mRNA libraries (∼20 million paired-end reads per sample) were generated on the NovaSeq X Plus platform. Reads were quantified with Salmon [29] using a decoy-aware index built from the *Ovis aries* reference genome (ARS-UI_Ramb_v3.0, RefSeq GCF_016772045.2) and transcriptome FASTA. The gentrome was created by concatenating genome and transcriptome FASTAs, with genome identifiers forming the decoy list. Salmon was run with --validateMappings and --gcBias, producing transcript-level TPMs and counts. Transcript quantifications were imported into R via tximport and summarized to gene-level count matrices. Genes with average TPM < 1 were filtered out. Normalization and differential expression were performed with DESeq2 [30], applying Benjamini–Hochberg correction; genes with adjusted p < 0.05 were considered significant. Functional enrichment (GO and pathways) was conducted using gProfiler2 and clusterProfiler, and visualizations (PCA plots, heatmaps, dot plots) were generated in R with ggplot2, pheatmap, and related packages.

### Lentiviral Production and infection

Production of lentiviral particles was performed as previously described [31] by polyethyleneimine mediated transfection of 293T cells with expression and packaging plasmids (psPAX2 and pMD2.G encoding Gag, Pol, and Rev). The following expression vectors were used PL-SIN-EOS-S(4+)-EiP (Addgene plasmid # 21314) and PL-SIN-EOS-C(3+)-EiP (Addgene plasmid # 21313) [12,13]. Viral supernatants were collected at 48 and 72 h, pooled and filtered using 0.45 µm PES filters. Viruses were concentrated by ultracentrifugation at 21,000 g for 90 min. Transduction was performed by adding concentrated lentiviruses to the culture medium and incubating for 24 h. Control GFP vector, pLX_TRC209 were used to estimate infection rates associated with viral preparation batches. pLX_TRC209 was a gift from David Root (Addgene plasmid # 125715). To differentiate the reporter lines, the cells were cultured in cell culture medium supplemented with 20% FBS. To evaluate the role of TGF-β inhibition on the reporter lines, 1μM A83-01 was added to the media. Fluorescence was measured using ImageJ software from the images captured under same exposure and gain conditions using Leica fluorescent microscope.

### Quantitative Real-Time PCR Analysis

RNA was extracted from the cells using RNeasy Plus Micro Kit (Qiagen). Complementary DNA (cDNA) was made using High-Capacity RNA-to-cDNA™ Kit (Applied biosystems). The primers used were GAPDH: F-5’GTTCCACGGCACAGTCAAGG-3’ and R-5’ACTCAGCACCAGCATCACCC-3’; OCT4: F-5’AACGAGAATCTGCAGGAGATATG-3’ and R- 5’TCTCACTCGGTTCTCGATACT-3’. PowerTrack™ SYBR Green Master Mix (Applied Biosystems) was used for qPCR as per manufacturer’s instructions. All reactions were run in triplicate. GAPDH served as the endogenous control. Relative expression levels were calculated using the ΔΔCt method, and results are presented as fold change (mean ± SEM). Statistical comparisons between control and treatment groups were performed using Student’s t-test using Prism.

## Results

### De novo derivation and characterization of sheep embryonic stem cells (sESCs) from in vivo blastocysts

Zona pellucida-free, day 7 in vivo-derived sheep blastocysts were plated onto mitomycin C-inactivated mouse embryonic fibroblasts (iMEFs) in a chemically defined N2B27-based medium supplemented with the WNT pathway inhibitor IWR1, together with Activin A and FGF2 (NBFR medium). Initial attachment was slow, and outgrowths of trophectoderm-like cells predominated (Fig. 1A; Supplementary Fig. 1A–B). However, with continued passaging, compact monolayer colonies with distinct borders subsequently emerged. From three blastocysts, two independent lines were established and stably maintained. These lines propagated as undifferentiated, flat monolayer colonies composed of tightly packed small polygonal cells with multiple prominent nucleoli (Fig 1A). Immunocytochemistry confirmed expression of cell surface pluripotency marker alkaline phosphatase (ALP) as well as core pluripotency markers OCT4/POU5F1 and SOX2 (Fig 1B). Both lines proliferated rapidly, with a doubling time of approximately 15 hours, and could be reliably passaged as single cells (Fig. 1C). Transcripts of trophectoderm and primitive endoderm–associated genes were absent or markedly reduced, whereas pluripotency-associated genes including OCT4, LIN28A, SALL4, and DNMT3B were highly expressed. Colonies also expressed high levels of primed pluripotency markers including DUSP6, GLI2, and ETV5, confirming their primed pluripotent state (Fig 1D,E). Karyotypic analysis confirmed genomic stability (Fig 1F). Immunostaining for the chromatin modification histone H3 lysine 27 trimethylation (H3K27me3) revealed a single, intense nuclear focus, consistent with X-chromosome inactivation in these female lines (Fig 1G). When cultured as single-cell suspensions in serum-containing medium devoid of growth factors and small molecule inhibitors, sESCs readily aggregated to form compact, spherical embryoid bodies within 48–72 hours. Upon subsequent attachment to gelatin-coated surfaces, these embryoid bodies produced adherent outgrowths displaying diverse morphologies characteristic of spontaneous differentiation. Immunocytochemical analysis of these differentiated outgrowths confirmed expression of lineage-associated markers indicative of differentiation into different lineages (Fig 1H). Furthermore, subcutaneous injections of these cells suspended in Matrigel® into immunodeficient NSG mice resulted in robust teratoma formation within six weeks. Histological examination confirmed tissue diversity, including neural rosettes and squamous epithelium (ectoderm), glandular structures (endoderm), and cartilage (mesoderm), confirming the formation of derivatives from all three embryonic germ layers (Fig 1H). The colonies also expanded robustly in an enhanced mTeSR medium (emTeSR; mTeSR supplemented with Activin A, IWR1, and FGF2), maintaining their morphology and pluripotency without the need for low–fatty acid BSA supplementation (Fig 1I). Under both NBFR and emTeSR conditions, flat colonies grew on iMEF feeders with sharply defined edges, resembling human primed embryonic stem cells, and could be enzymatically dissociated and passaged as single cells every 2–3 days at a 1:4 ratio. Based on these morphological and molecular features, the established lines were designated sheep embryonic stem cells (sESCs).

**Figure 1.**
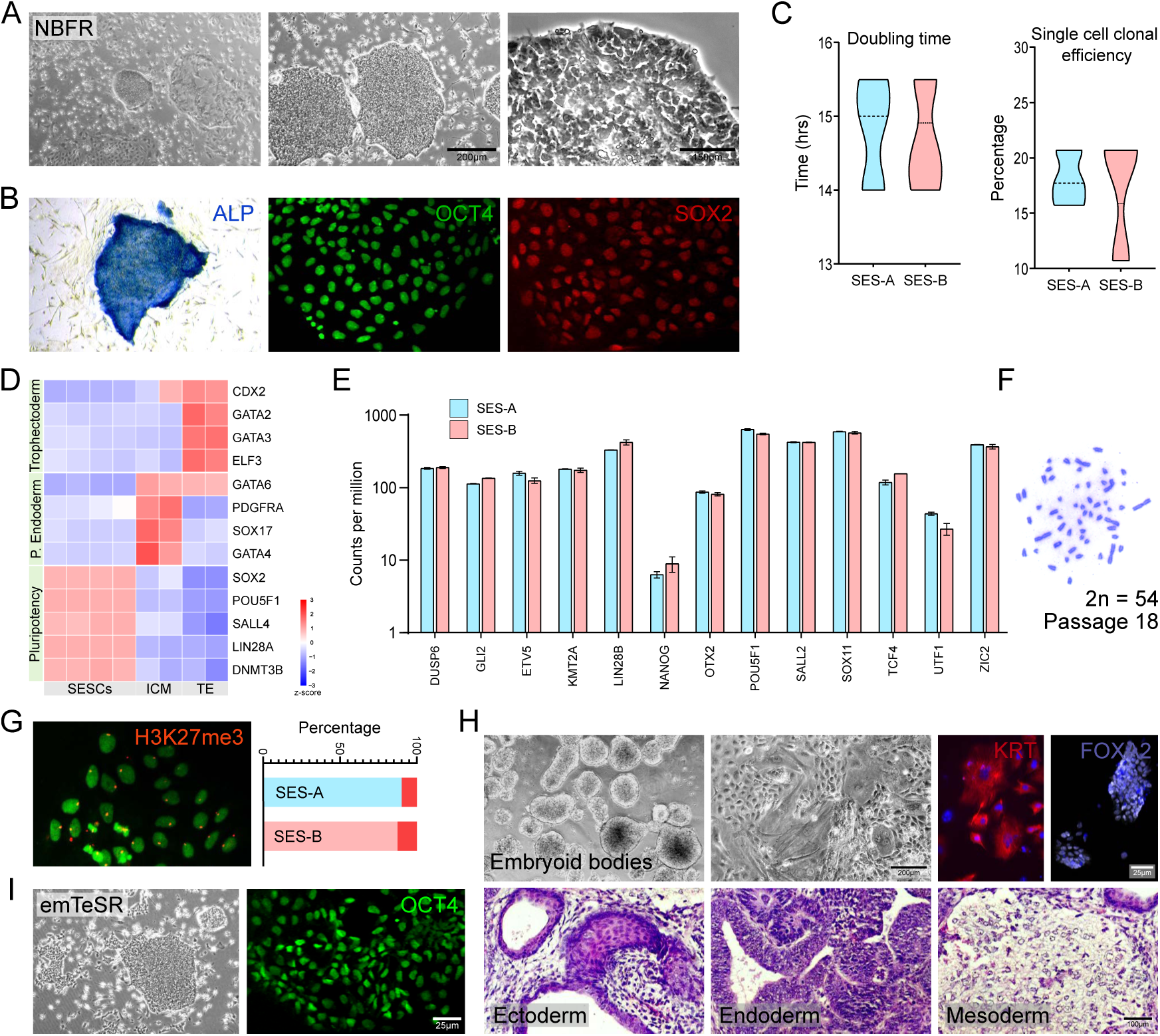
Derivation, characterization, and maintenance of sheep embryonic stem cell (sESC) lines. (A) Primed sESC lines were derived from in vivo blastocyst in NBFR medium formed compact monolayer colonies with sharp borders. (B) Colonies expressed pluripotency markers ALP, OCT4/POU5F1, and SOX2. (C) Lines proliferated rapidly (doubling time ∼15 h) and were efficiently passaged as single cells. (D–E) Pluripotency-associated genes and primed markers were highly expressed, with trophectoderm and primitive endoderm genes absent or markedly reduced. (F) Karyotypes were normal. (G) H3K27me3 immunostaining showed a single nuclear focus, consistent with X-chromosome inactivation. (H) sESCs formed compact embryoid bodies that differentiated into cells expressing lineage markers, and produced teratomas containing ectodermal, mesodermal, and endodermal derivatives in NSG mice. (I) Cells also expanded robustly in emTeSR medium (mTeSR + Activin A, IWR1, FGF2), maintaining primed ESC-like morphology and OCT4 expression.

### Feeder free culture of sESCs

To assess the potential for feeder-free propagation of sESCs, we evaluated a panel of commonly used extracellular matrix substrates, including collagen, laminin, Geltrex, Cultrex, Matrigel, and vitronectin. In NBFR medium, sESCs failed to attach to collagen coated dishes and exhibited poor attachment on laminin. Although colonies initially expressed OCT4 and SOX2 on laminin-coated surfaces, cells frequently rounded up, detached, and failed to establish long-term cultures. In contrast, robust adhesion and proliferation were observed on Geltrex, Cultrex, Matrigel, and vitronectin, and the colonies were alkaline phosphatase positive (Fig. 2, Supplementary Fig. 2A). On Geltrex, Cultrex, and Matrigel, sESCs grew as monolayers with indistinct colony borders, whereas colonies on vitronectin displayed sharper boundaries. However, when cultured in emTeSR medium, colonies on Geltrex, Cultrex, Matrigel, and vitronectin exhibited well-defined borders, retained alkaline phosphatase activity, and maintained strong OCT4 and SOX2 expression (Fig. 2, Supplementary Fig. 2A). Under feeder-free conditions, sESCs could be stably maintained long term with homogeneous morphology and sustained expression of pluripotency markers. The WNT pathway inhibitor IWR1 could be replaced by IWP2 or XAV939, both of which supported colonies with compact, sharply demarcated edges over multiple passages. However, omission of WNT inhibition led to differentiation, evidenced by the loss of colony definition and the emergence of morphologically distinct differentiated cells (Supplementary Fig. 2B). Collectively, these findings indicate that colony morphology is influenced by both the extracellular matrix substrate and the culture medium.

**Figure 2.**
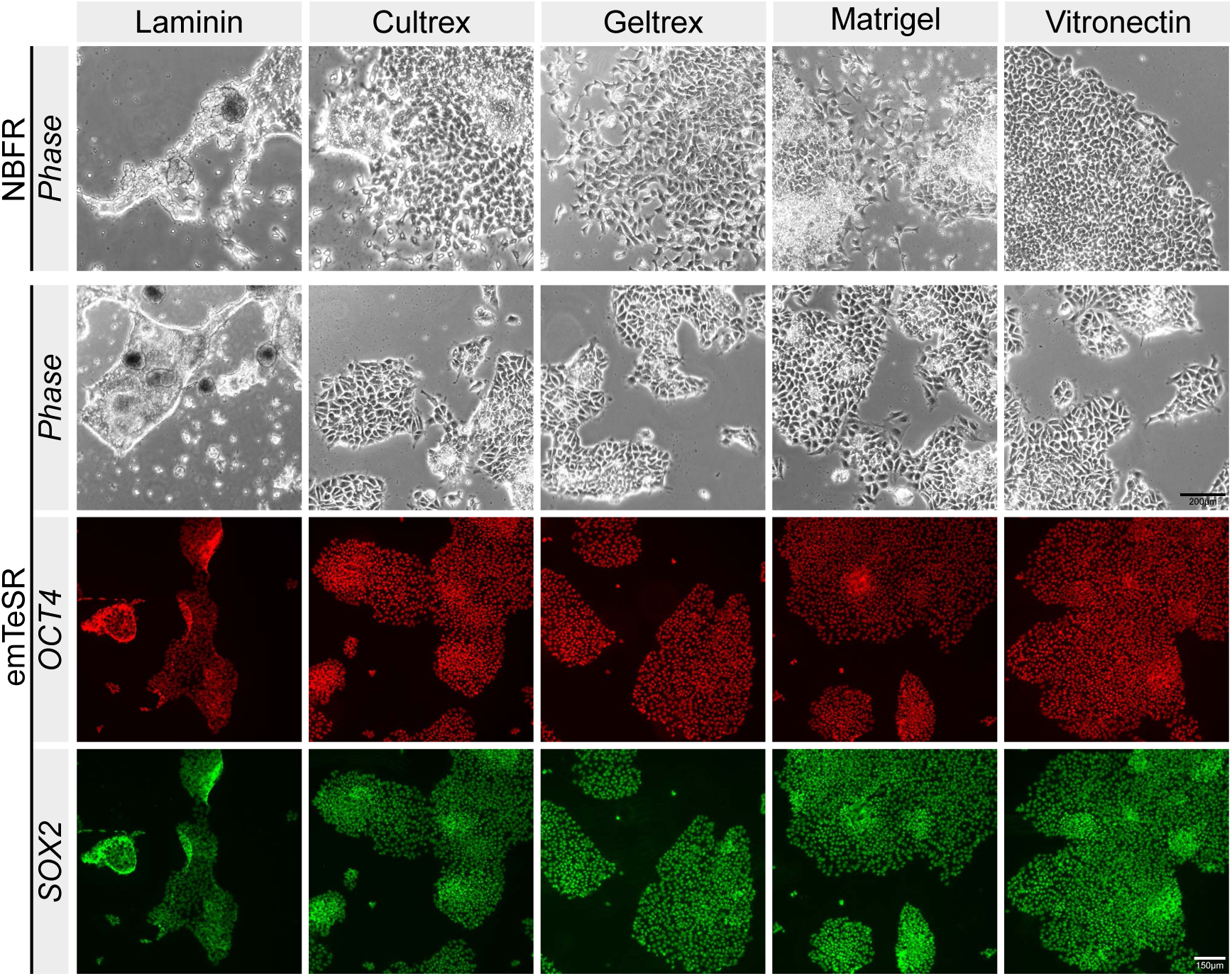
Feeder free propagation of sESCs. Feeder-free culture conditions were assessed on collagen, laminin, Geltrex, Cultrex, Matrigel, and vitronectin. In NBFR medium, sESCs failed to attach on collagen and showed poor, unstable attachment on laminin. Geltrex, Cultrex, Matrigel, and vitronectin supported robust adhesion and proliferation, with indistinct colony borders on Geltrex, Cultrex, and Matrigel and sharper borders on vitronectin. In emTeSR medium, all four substrates supported well-defined colonies that maintained strong OCT4 and SOX2 expression.

### OCT4- and SOX2-enhancer–driven GFP reporters robustly mark pluripotent sESCs

To evaluate whether enhancer-driven reporters faithfully identify pluripotent sESCs, we transduced sESCs with EOS-lentiviral vectors containing either the multimerized Oct4 core enhancer element CR4 (conserved region 4) or the Sox2 core enhancer element SRR2 (Sox2 regulatory region 2). These elements were coupled to the early transposon promoter driving an EGFP-IRES-PuroR cassette, previously shown to reliably mark pluripotent stem cells in mouse and human models [12,13]. Following puromycin selection, surviving sESC colonies maintained strong and stable EGFP expression (Fig. 3A). Comparable GFP expression was observed in cells infected with a vector in which GFP was driven by a constitutively active EF1α promoter (Fig. 3B). Importantly, lentiviral infection did not alter endogenous OCT4 or SOX2 expression in EGFP-positive lines (Fig. 3C–D), regardless of whether cells were infected with EOS or EF1α vectors. However, upon in vitro differentiation, EGFP expression was extinguished in EOS-infected sESCs in parallel with loss of Oct4, whereas EF1α-GFP lines maintained GFP despite losing OCT4 expression (Fig. 3E–I). This divergence confirms that enhancer-driven GFP reporters selectively reflect the pluripotent state, while constitutive reporters do not. Consistent with the linked IRES-PuroR design, differentiated EOS-infected cells also lost puromycin resistance (data not shown). To further validate the responsiveness of EOS reporters, we examined their ability to detect the known dependence of primed pluripotency on Activin/TGFβ signaling. Pharmacologic inhibition of Activin/TGFβ receptors with A83-01 induced differentiation and resulted in loss of GFP in EOS-OCT4-GFP cells, accompanied by downregulation of Oct4 transcripts (Fig. 4J–L). These findings establish EOS-reporter sESC lines as specific live-cell tools for tracking and monitoring pluripotency in real time, providing valuable resources for pluripotency studies.

**Figure 3.**
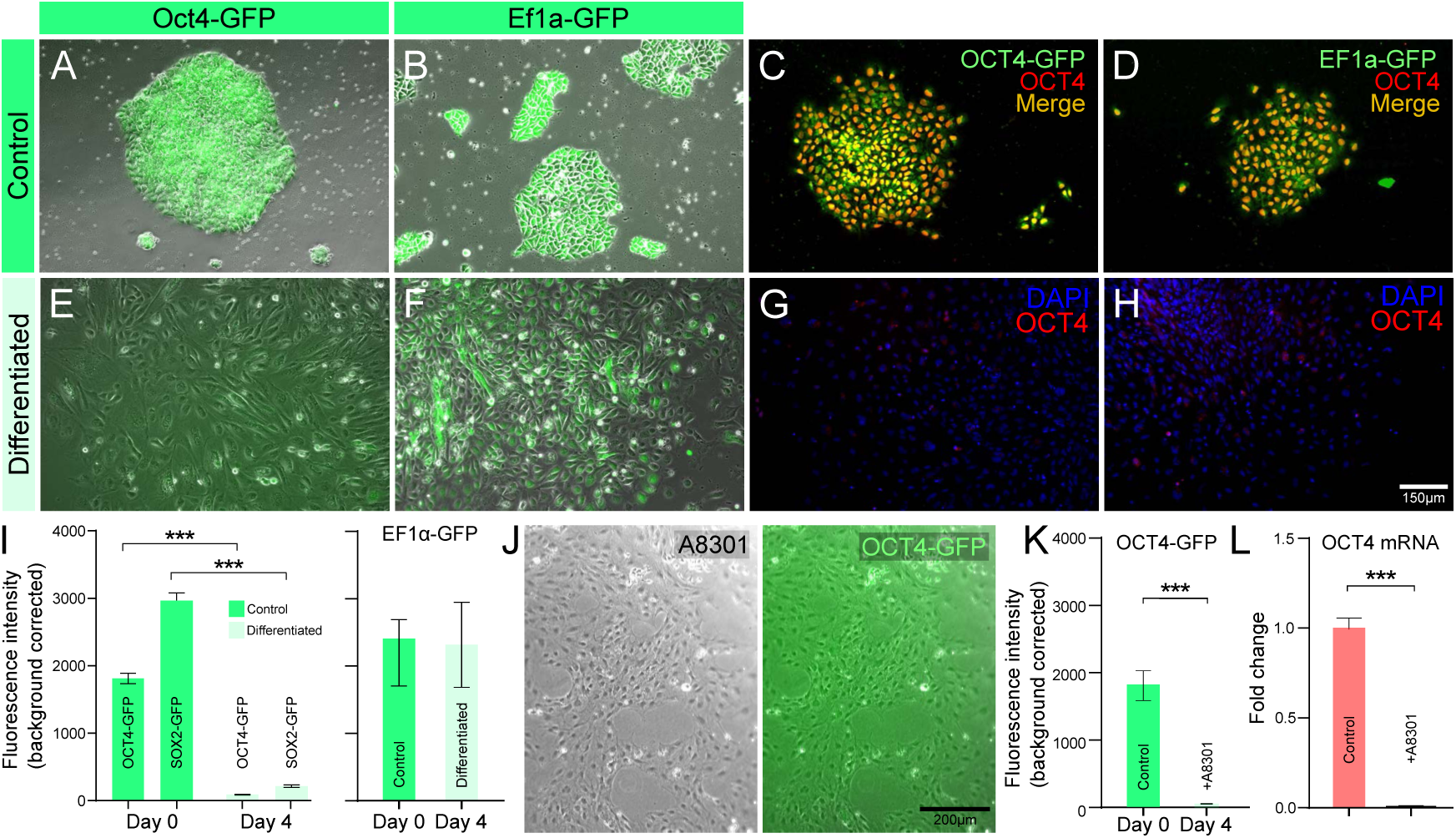
OCT4- and SOX2–enhancer–driven GFP reporters selectively mark pluripotent sESCs. (A,B) Undifferentiated sESC colonies infected with EOS vectors containing the OCT4 core enhancer CR4 driving GFP, as well as EF1α-GFP controls, showed strong and stable EGFP expression. (C,D) Lentiviral transduction did not alter endogenous OCT4 expression in EGFP-positive cells. (E–H) Upon in vitro differentiation, EGFP expression was extinguished in EOS-infected cells in parallel with loss of Oct4, whereas EF1α-GFP expression persisted despite Oct4 downregulation, confirming enhancer specificity for the pluripotent state. (I) Fluorescence quantification further demonstrated loss of GFP intensity in EOS-Oct4-GFP and EOS-Sox2-GFP cells following differentiation, while GFP remained stable in EF1α-GFP controls. (J–L) Inhibition of Activin/TGFβ signaling with A83-01 induced differentiation and resulted in loss of GFP in EOS-Oct4-GFP lines, accompanied by reduced Oct4 transcript levels.

**Figure 4.**
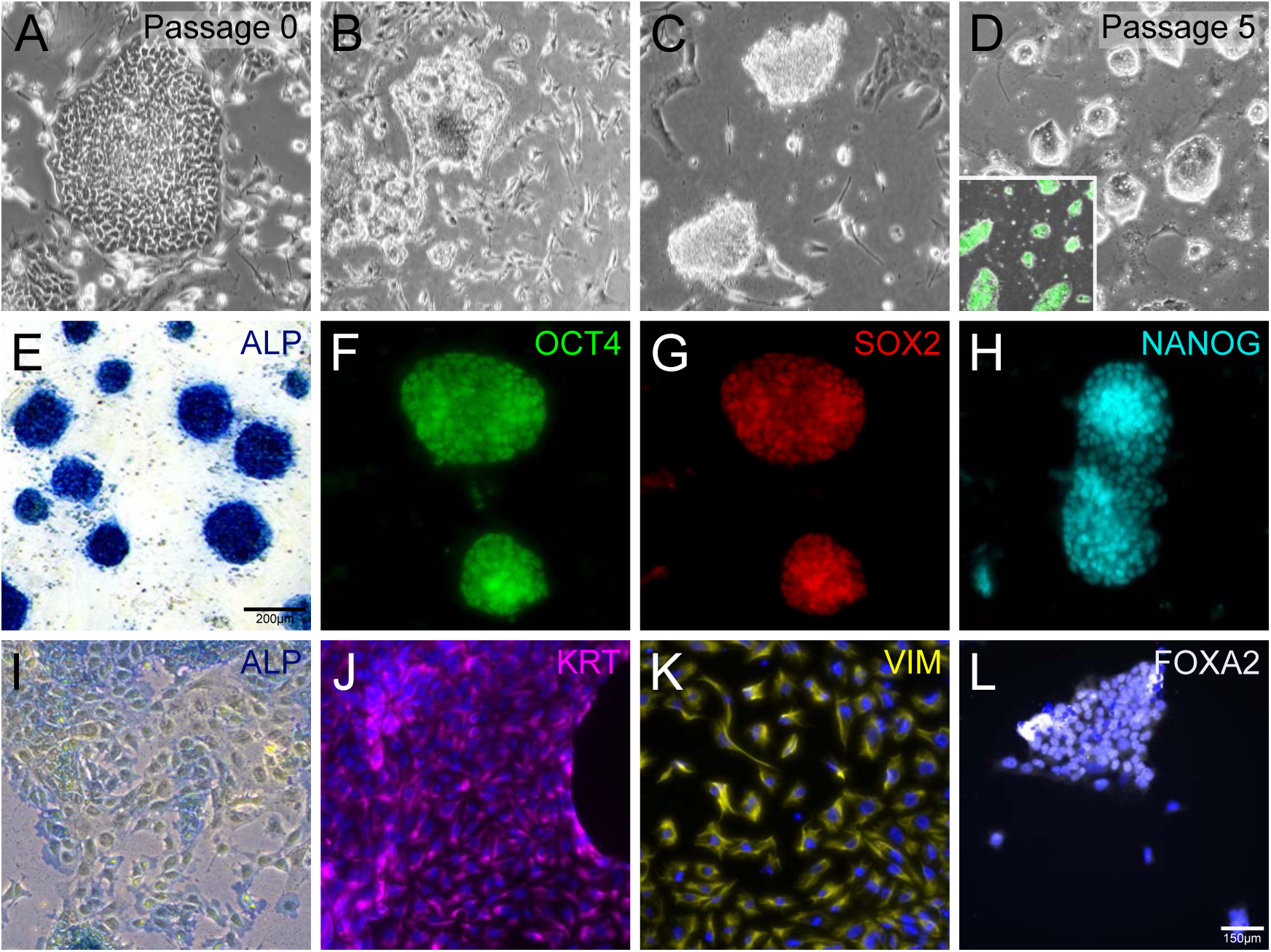
Conversion of sESCs into an alternate pluripotent state. (A–D) Flattened sESC EOS-OCT4-GFP cultures transferred into HENSM medium initially underwent substantial cell death, but by passages 3–5, compact EGFP-positive dome-shaped colonies emerged. (E–H) Dome-shaped colonies were independently generated from both primed lines (sESa and sESb), retained alkaline phosphatase staining, and expressed OCT4, SOX2, and NANOG. (I-L) Upon exposure to serum-containing differentiation medium, colonies flattened, lost ALP activity, and produced cells expressing markers of all three germ layers.

### Conversion of sESCs into an alternate pluripotent state

Recent studies in mice and humans have identified culture conditions capable of reverting primed cells to a naïve state [32]. To test whether primed sESCs could similarly be converted, we examined several naïve culture conditions developed for mouse and human ESCs, including 2i/LIF [24], t2iL+Gö [25], 5i/L/A [23], and the recently reported HENSM medium [33]. EOS-OCT4-GFP colonies cultured in 2i/LIF, t2iL+Gö, or 5i/L/A either failed to survive or lost EGFP expression and alkaline phosphatase positivity, indicating differentiation (Supplementary Fig. 4 A,B). In contrast, when cultured in HENSM medium, widespread cell death occurred during the initial two days of adaptation; however, by passage 3-5, a subset of compact, EGFP positive dome-shaped colonies emerged (Fig. 4A–D). These colonies could be stably propagated in HENSM medium for at least 20 passages without detectable differentiation, retaining uniform dome-shaped morphology and normal karyotypes (Supplementary Fig. 3C). Similar dome-shaped colonies were independently derived from both primed lines (sESa and sESb). The established colonies were alkaline phosphatase–positive and expressed key pluripotency markers, including OCT4, SOX2, and NANOG (Fig. 4E–H). When exposed to serum-containing differentiation medium, the colonies flattened, lost alkaline phosphatase activity, and gave rise to cells expressing markers representative of the three embryonic germ layers, confirming their trilineage differentiation potential (Fig. 4I–L). Although these cells could be maintained under both feeder-dependent and feeder-free conditions, cultures displayed more uniform morphology and growth in the presence of feeders. The converted lines were readily passaged using TrypLE every 4–5 days and maintained robust proliferation. Collectively, these findings indicate that sESCs can be reprogrammed under HENSM conditions into a stable, alternate pluripotent state that is morphologically and functionally distinct from the primed state.

### Sheep intermediate pluripotent stem cells exhibit molecular and functional features closer to a naïve pluripotent state

To further investigate the molecular characteristics of the sheep intermediate pluripotent stem cells, we performed RNA-sequencing (RNA-seq) on both the intermediate cells and their parental sESCs. Pearson correlation analysis showed that the intermediate cells formed a distinct transcriptional group, clearly separated from the primed sESCs derived in this study and from previously reported primed sESCs [19] (Figure 5A). To further resolve the gene regulatory networks underlying these differences, we performed an integrated analysis that included parental primed sESCs, previously published bovine ESCs (bESC) [17], human primed ESCs (hPrimed) [34], human expanded potential stem cells (hEPSC) [35], human naïve ESCs (hNaïve) [34], and sheep blastocyst inner cell mass (ICM) cells [19]. Principal component analysis (PCA) revealed that the sheep intermediate pluripotent stem cells clustered as a discrete group, separating prominently along PC2. Notably, they occupied a position closer to human naïve ESCs and sheep ICM cells along PC1, indicating a transcriptional landscape more closely aligned with naïve than with primed pluripotency (Fig. 5B). Gene enrichment analysis showed that, relative to parental sESCs, the intermediate cells significantly upregulated developmental and pluripotency-associated pathways, particularly those involving matrix interactions and signaling regulation (Fig. 5C). At the level of individual transcripts, the intermediate cells increased expression of key naïve-associated markers, including NANOG, STAT3, TFCP2L1, KLF4, KLF5, ELF3, and FOXI3 (Figure 5D–E). TFCP2L1, a critical regulator required for resetting primed cells toward a naïve state [36], was prominently upregulated. In addition to naïve-associated factors, transcripts characteristic of the formative phase of pluripotency, such as ZIC5, ZIC2, OTX2, and UTF1, were also highly upregulated. In contrast, genes associated with primed pluripotency and expanded potential states were downregulated.

**Figure 5.**
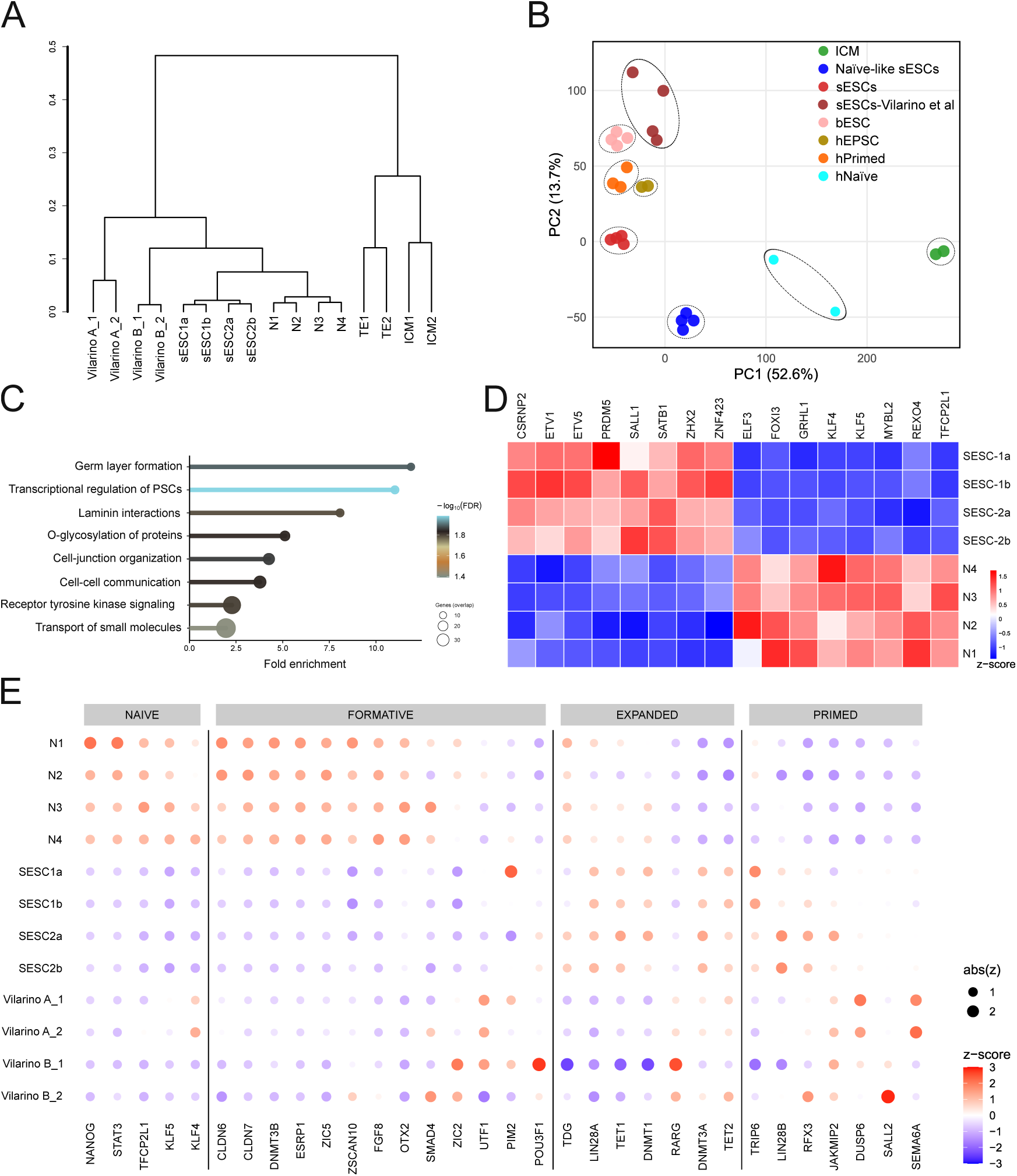
Transcriptional profiling of sESCs grown in HENSM medium. (A) Pearson correlation analysis shows that sESCs cultured in HENSM medium form a distinct transcriptional group, clearly separated from the primed sESCs generated in this study and previously reported primed psESCs. (B) Principal component analysis places HENSM-grown sESCs as a discrete population, separating prominently along PC2 and positioned closer to human naïve ESCs and sheep ICM cells along PC1. (C) Gene enrichment analysis highlighting pathways upregulated under HENSM conditions. (D) Heat map comparing primed sESCs and HENSM-grown sESCs, showing upregulation of naïve-associated genes and downregulation of primed markers. (E) Dot plots of representative pluripotency-state markers demonstrating that HENSM conditions convert sESCs into a transcriptionally distinct intermediate state characterized by increased expression of naïve and formative pluripotency markers.

## Discussion

In this study, we report the derivation and long-term maintenance of sheep embryonic stem cells (sESCs) from in vivo blastocysts under chemically defined conditions and demonstrate their conversion into a stable intermediate pluripotent state with naïve-like features. These findings address long-standing challenges in capturing authentic pluripotent stem cell states in ungulate species and provide new experimental platforms for ruminant stem cell biology, genetic engineering, and cellular agriculture.

Previous reports of ovine pluripotent stem cells have described sheep ESCs in CTFR medium [19] or embryonic disc stem cells in XAV-939–based systems [20], yet culture outcomes differed in morphology and stability. Here, we established primed sESCs under two distinct culture conditions, both on feeders and under feeder-free systems. A central observation in this study is the strong influence of medium composition on sESC morphology and stability. In NBFR medium, sESCs adopted flattened monolayer morphologies with indistinct colony borders, reminiscent of bovine ESCs and other ungulate pluripotent systems. By contrast, an enhanced mTeSR-based formulation (emTeSR), supplemented with Activin A, IWR1, and FGF2, supported compact colonies with sharply demarcated edges and robust expression of pluripotency markers. This qualitative improvement likely reflects the stabilized TGF-β/Activin signaling provided by the mTeSR basal formulation in combination with exogenous Activin A, consistent with the established requirement for TGF-β pathway activity in maintaining primed pluripotency in ungulate species [20,21] . Although the potential contribution of other medium components cannot be excluded, this explanation is supported by earlier findings that CTFR medium, based on the same basal formulation but lacking TGF-β or Activin, failed to sustain feeder-free growth [17,19]. Accordingly, pharmacologic inhibition of TGF-β signaling induced rapid differentiation, reinforcing its essential role in sESC maintenance. Together, these observations highlight the pivotal influence of medium composition on ungulate ESC derivation. While NBFR offers simplicity and defined composition, the emTeSR formulation provides enhanced morphological stability and molecular maintenance of pluripotency, representing a key advancement in culture systems for sheep primed pluripotent stem cells.

The successful transition of sESCs to feeder-free culture conditions across multiple extracellular matrix substrates represents a significant practical advancement that expands their applicability for both basic research and translational use. The substrate-dependent variation in colony morphology, ranging from indistinct monolayers on some basement membrane matrices to sharply delineated colonies on vitronectin, reflects the interplay between integrin signaling, cytoskeletal tension, and ECM biochemical composition in shaping pluripotent cell behavior [37–39]. Similar effects have been documented in human and mouse systems, where ECM components influence colony cohesion, pluripotency gene expression, and sensitivity to differentiation cues [39–41]. The ability to maintain sESCs with normal karyotypes and stable expression of pluripotency markers across compatible feeder-free conditions establishes a robust and scalable platform for genetic modification, disease modeling, and bioprocess development in sheep.

In parallel with culture optimization, we developed and validated EOS-lentiviral reporter lines driven by OCT4 (CR4) and SOX2 (SRR2) enhancer elements. These reporters faithfully marked pluripotent sESCs, were extinguished upon in vitro differentiation, and showed high sensitivity to perturbation of Activin/TGF-β signaling. Their behavior closely parallels that observed in mouse and human ESCs, where EOS reporters have proven invaluable for dissecting pluripotency networks and tracking state transitions [12,13]. In the ovine system, these reporter lines fill a critical gap by enabling real-time, live-cell monitoring of pluripotency and differentiation, and by providing a direct functional readout of signaling pathway dependencies. As such, they represent a versatile tool for future mechanistic studies and for optimizing conditions for reprogramming, differentiation, and interspecies chimera formation.

A major conceptual advance of this work is the demonstration that sESCs can be reconfigured from a primed state into a stable intermediate pluripotent state under defined HENSM conditions. Conventional mouse and human naïve media (2i/LIF, t2iL+Gö, 5i/L/A) were insufficient to support sESC survival or pluripotency, underscoring fundamental interspecies differences in signaling requirements for naïve state stabilization. In mouse, naïve ESCs rely on dual inhibition of MEK and GSK3 pathways alongside LIF/STAT3 activation [24]; however, these conditions failed to maintain OCT4 and led to differentiation in sheep ESCs. Even in humans, naïve-like states captured by 5i/L/A or related cocktails often display partial instability and require extensive pathway modulation [23,25]. By contrast, HENSM conditions supported the emergence of dome-shaped, EGFP-positive colonies with long-term stability, normal karyotype, and trilineage potential. The ability of these cells to expand over multiple passages indicates that sheep pluripotency networks can be reconfigured into a state distinct from both conventional primed and naïve states. Transcriptomic profiling revealed that these cells occupy a unique position along the pluripotency continuum. They expressed hallmark naïve-associated genes—including NANOG, STAT3, KLF4, KLF5, ELF3, and especially TFCP2L1, a master regulator required for resetting primed cells toward naïve identity [36]—while simultaneously upregulating formative-associated genes such as ZIC2, ZIC5, OTX2, and UTF1. This hybrid molecular signature suggests that these cells do not represent a canonical “ground state” but instead resemble an intermediate or formative-like configuration. Similar states have been described in rodent, bovine, and human systems, where transitional pluripotent cells retain developmental competence but exhibit partial activation of lineage-priming programs [18,35,42]. Our PCA analysis further distinguished sheep intermediate cells from human EPSCs, indicating that they represent a distinct, species-specific regulatory state rather than a direct analog of expanded potential pluripotency.

These findings are in line with emerging models that view pluripotency not as a simple binary switch between naïve and primed states but as a continuum encompassing transitional and region-specific states [43–45]. Our findings also align with prior evidence from cattle and pigs, where attempts to capture naïve pluripotency often yield unstable or partially reset states [16,18,46]. The long-term stability, normal karyotypes, and robust differentiation potential of sheep intermediate cells suggest that this state may represent an optimal equilibrium between developmental potency and culture stability in ruminant species. However, deeper characterization will be essential to define their precise position along the pluripotency continuum. Future studies should include genome-wide DNA methylation profiling, assessment of enhancer usage, mapping of chromatin accessibility, and evaluation of X-chromosome activation status in female lines. The definitive functional test of naïve pluripotency, robust contribution to chimerism and germline transmission following blastocyst injection, will be crucial to establishing whether these sheep cells can achieve bona fide naïve potential.

In conclusion, this study establishes a robust framework for sheep pluripotent stem cell research. We have developed a robust culture platform for primed sESCs, generated validated reporter lines, and, most importantly, described the capture of a stable naïve-like intermediate cell type. These resources provide new tools for investigating ungulate development, creating advanced models for genetic engineering in agriculture, and deepening our comparative understanding of the fundamental principles of pluripotency across mammals.

## Acknowledgements

We thank Gerald R. Kelly at the Purdue University Animal Sciences Research and Education Center (ASREC) Sheep unit for breeding the ewes for embryo flushing.

## Author contributions

VVP and MAV conceived the study and designed the experiments. STS performed cell line derivation, molecular characterization, transcriptome data analysis, immunocytochemistry, and imaging. VVP and STS analyzed the data and interpreted the results. STS and VVP wrote the initial draft of the manuscript, and all authors contributed to editing and approved the final version. VVP supervised the project.

## Data availability

The RNA-seq datasets generated and analyzed during this study have been deposited in the NCBI Gene Expression Omnibus (GEO) under accession number GSE311245. All other relevant data supporting the findings of this study are included in the manuscript and supplementary material.

---

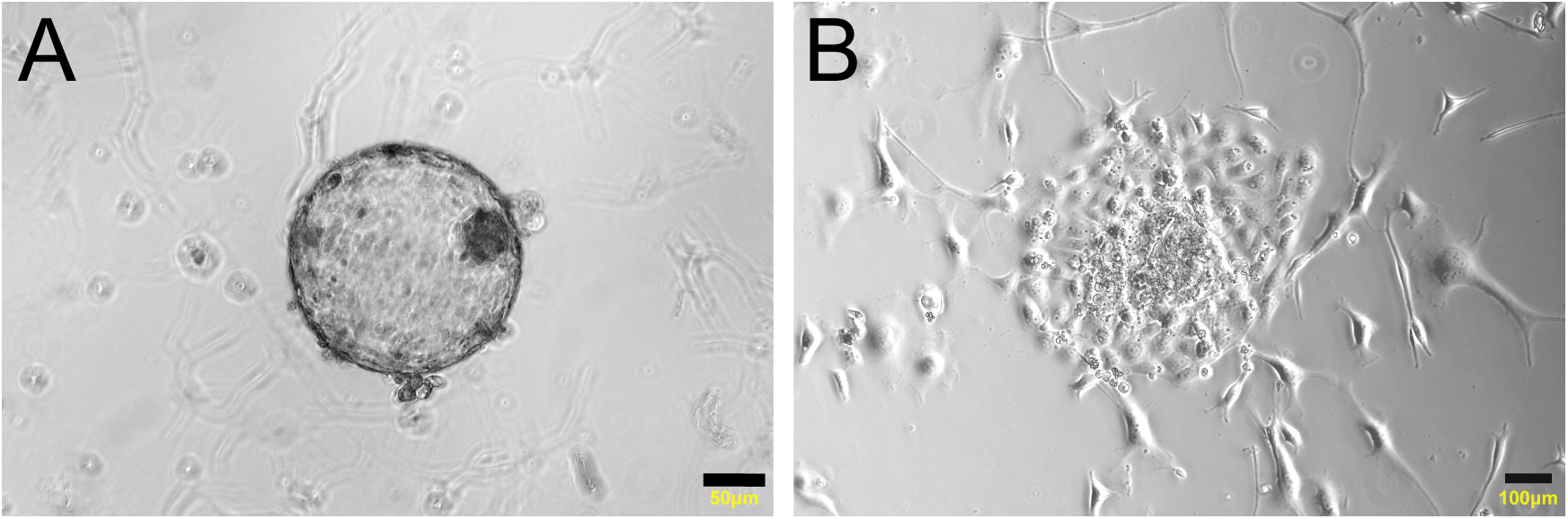

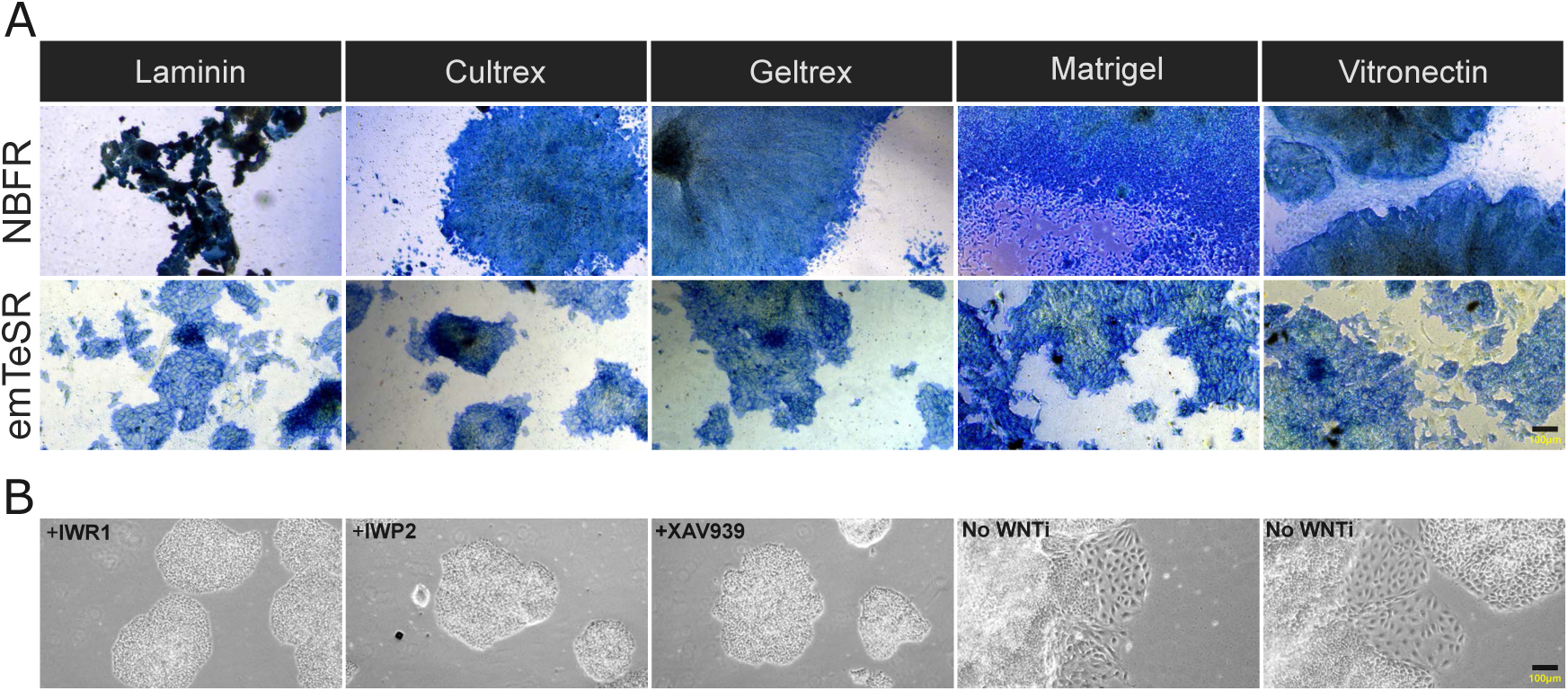

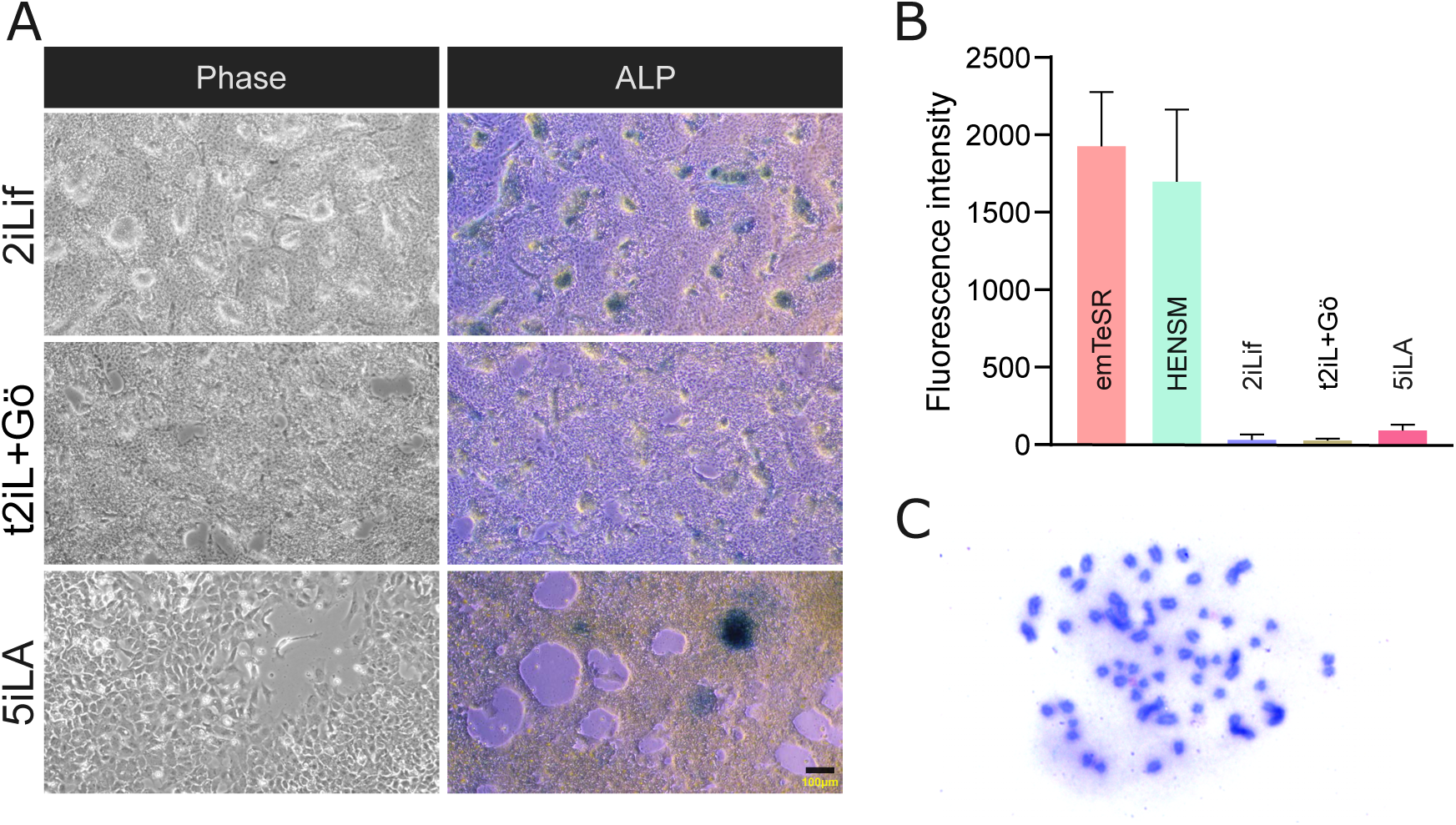

## References

[1] Evans MJ, Kaufman MH. Establishment in culture of pluripotential cells from mouse embryos. Nature 1981; 292:154–156.

[2] Lee DK, Kim M, Jeong J, Lee YS, Yoon JW, An MJ, Jung HY, Kim CH, Ahn Y, Choi KH, Jo C, Lee CK. Unlocking the potential of stem cells: Their crucial role in the production of cultivated meat. Curr Res Food Sci 2023; 7:100551.

[3] Navarro M, Laiz-Quiroga L, Blüguermann C, Mutto A. Livestock embryonic stem cells for reproductive biotechniques and genetic improvement. Anim Reprod 2024; 21:e20240029.

[4] Singh P, Ali SA. Impact of CRISPR-Cas9-Based Genome Engineering in Farm Animals. Vet Sci 2021; 8:122.

[5] Laustriat D, Gide J, Peschanski M. Human pluripotent stem cells in drug discovery and predictive toxicology. Biochem Soc Trans 2010; 38:1051–1057.

[6] Dundes CE, Loh KM. Bridging naïve and primed pluripotency. Nat Cell Biol 2020; 22:513–515.

[7] Kinoshita M, Barber M, Mansfield W, Cui Y, Spindlow D, Stirparo GG, Dietmann S, Nichols J, Smith A. Capture of Mouse and Human Stem Cells with Features of Formative Pluripotency. Cell Stem Cell 2021; 28:453–471.e8.

[8] Yu L, Wei Y, Sun HX, Mahdi AK, Pinzon Arteaga CA, Sakurai M, Schmitz DA, Zheng C, Ballard ED, Li J, Tanaka N, Kohara A, et al. Derivation of Intermediate Pluripotent Stem Cells Amenable to Primordial Germ Cell Specification. Cell Stem Cell 2021; 28:550–567.e12.

[9] Lackner A, Sehlke R, Garmhausen M, Stirparo GG, Huth M, Titz-Teixeira F, Lelij P van der, Ramesmayer J, Thomas HF, Ralser M, Santini L, Galimberti E, et al. Cooperative genetic networks drive embryonic stem cell transition from naïve to formative pluripotency. EMBO J 2021; 40.

[10] Wang X, Wu Q. The Divergent Pluripotent States in Mouse and Human Cells. Genes (Basel) 2022; 13:1459.

[11] Neagu A, van Genderen E, Escudero I, Verwegen L, Kurek D, Lehmann J, Stel J, Dirks RAM, van Mierlo G, Maas A, Eleveld C, Ge Y, et al. In vitro capture and characterization of embryonic rosette-stage pluripotency between naive and primed states. Nat Cell Biol 2020; 22:534–545.

[12] Hotta A, Cheung AYL, Farra N, Vijayaragavan K, Séguin CA, Draper JS, Pasceri P, Maksakova IA, Mager DL, Rossant J, Bhatia M, Ellis J. Isolation of human iPS cells using EOS lentiviral vectors to select for pluripotency. Nat Methods 2009; 6:370–376.

[13] Hotta A, Cheung AYL, Farra N, Garcha K, Chang WY, Pasceri P, Stanford WL, Ellis J. EOS lentiviral vector selection system for human induced pluripotent stem cells. Nat Protoc 2009; 4:1828–1844.

[14] Pillai V V., Kei TG, Reddy SE, Das M, Abratte C, Cheong SH, Selvaraj V. Induced pluripotent stem cell generation from bovine somatic cells indicates unmet needs for pluripotency sustenance. Animal Science Journal 2019; 90:1149–1160.

[15] Ruan D, Xuan Y, Tam TTKK, Li ZX, Wang X, Xu S, Herrmann D, Niemann H, Lai L, Gao X, Nowak-Imialek M, Liu P. An optimized culture system for efficient derivation of porcine expanded potential stem cells from preimplantation embryos and by reprogramming somatic cells. Nat Protoc 2024; 19:1710–1749.

[16] Gao X, Nowak-Imialek M, Chen X, Chen D, Herrmann D, Ruan D, Chen ACH, Eckersley-Maslin MA, Ahmad S, Lee YL, Kobayashi T, Ryan D, et al. Establishment of Porcine and Human Expanded Potential Stem Cells. Nat Cell Biol 2019; 21:687.

[17] Bogliotti YS, Wu J, Vilarino M, Okamura D, Soto DA, Zhong C, Sakurai M, Sampaio RV, Suzuki K, Izpisua Belmonte JC, Ross PJ. Efficient derivation of stable primed pluripotent embryonic stem cells from bovine blastocysts. Proceedings of the National Academy of Sciences 2018; 115:2090–2095.

[18] Zhao L, Gao X, Zheng Y, Wang Z, Zhao G, Ren J, Zhang J, Wu J, Wu B, Chen Y, Sun W, Li Y, et al. Establishment of bovine expanded potential stem cells. Proceedings of the National Academy of Sciences 2021; 118:e2018505118.

[19] Vilarino M, Soto DA, Bogliotti YS, Yu L, Zhang Y, Wang C, Paulson E, Zhong C, Jin M, Belmonte JCI, Wu J, Ross PJ. Derivation of sheep embryonic stem cells under optimized conditions. Reproduction 2020; 160:761–772.

[20] Kinoshita M, Kobayashi T, Planells B, Klisch D, Spindlow D, Masaki H, Bornelöv S, Stirparo GG, Matsunari H, Uchikura A, Lamas-Toranzo I, Nichols J, et al. Pluripotent stem cells related to embryonic disc exhibit common self-renewal requirements in diverse livestock species. Development 2021; 148:dev199901.

[21] Soto DA, Navarro M, Zheng C, Halstead MM, Zhou C, Guiltinan C, Wu J, Ross PJ. Simplification of culture conditions and feeder-free expansion of bovine embryonic stem cells. Sci Rep 2021; 11.

[22] Bayerl J, Ayyash M, Shani T, Manor YS, Gafni O, Massarwa R, Kalma Y, Aguilera-Castrejon A, Zerbib M, Amir H, Sheban D, Geula S, et al. Principles of signaling pathway modulation for enhancing human naive pluripotency induction. Cell Stem Cell 2021; 28:1549–1565.e12.

[23] Theunissen TW, Powell BE, Wang H, Mitalipova M, Faddah DA, Reddy J, Fan ZP, Maetzel D, Ganz K, Shi L, Lungjangwa T, Imsoonthornruksa S, et al. Systematic identification of culture conditions for induction and maintenance of naive human pluripotency. Cell Stem Cell 2014; 15:471–487.

[24] Ying QL, Wray J, Nichols J, Batlle-Morera L, Doble B, Woodgett J, Cohen P, Smith A. The ground state of embryonic stem cell self-renewal. Nature 2008; 453:519–523.

[25] Takashima Y, Guo G, Loos R, Nichols J, Ficz G, Krueger F, Oxley D, Santos F, Clarke J, Mansfield W, Reik W, Bertone P, et al. Resetting Transcription Factor Control Circuitry toward Ground-State Pluripotency in Human. Cell 2014; 158:1254.

[26] Pillai VV, Siqueira LG, Das M, Kei TG, Tu LN, Herren AW, Phinney BS, Cheong SH, Hansen PJ, Selvaraj V. Physiological profile of undifferentiated bovine blastocyst-derived trophoblasts. Biol Open 2019; 8:bio037937.

[27] Pillai VV, Kei TG, Gurung S, Das M, Siqueira LGB, Cheong SH, Hansen PJ, Selvaraj V. RhoA/ROCK signaling antagonizes bovine trophoblast stem cell self-renewal and regulates preimplantation embryo size and differentiation. Development (Cambridge) 2022; 149.

[28] Pillai VV, Koganti PP, Kei TG, Gurung S, Butler WR, Selvaraj V. Efficient induction and sustenance of pluripotent stem cells from bovine somatic cells. Biol Open 2021; 10.

[29] Patro R, Duggal G, Love MI, Irizarry RA, Kingsford C. Salmon: fast and bias-aware quantification of transcript expression using dual-phase inference. Nat Methods 2017; 14:417.

[30] Love MI, Huber W, Anders S. Moderated estimation of fold change and dispersion for RNA-seq data with DESeq2. Genome Biology 2014 15:12 2014; 15:550-.

[31] Vijayan Pillai V, Koganti PP, Gurung S, Cheong SH, Selvaraj V. Transformed bovine trophoblast stem cell lines, characterization, gene editing and secretions. Biol Reprod 2025.

[32] Zhou J, Hu J, Wang Y, Gao S. Induction and application of human naive pluripotency. Cell Rep 2023; 42:112379.

[33] Bayerl J, Ayyash M, Shani T, Manor YS, Gafni O, Massarwa R, Kalma Y, Aguilera-Castrejon A, Zerbib M, Amir H, Sheban D, Geula S, et al. Principles of signaling pathway modulation for enhancing human naive pluripotency induction. Cell Stem Cell 2021; 28:1549–1565.e12.

[34] Hu Z, Li H, Jiang H, Ren Y, Yu X, Qiu J, Stablewski AB, Zhang B, Buck MJ, Feng J. Transient inhibition of mTOR in human pluripotent stem cells enables robust formation of mouse-human chimeric embryos. Sci Adv 2020; 6.

[35] Yang Y, Liu B, Xu J, Wang J, Wu J, Shi C, Xu Y, Dong J, Wang C, Lai W, Zhu J, Xiong L, et al. Derivation of Pluripotent Stem Cells with In Vivo Embryonic and Extraembryonic Potency. Cell 2017; 169:243–257.e25.

[36] Takashima Y, Guo G, Loos R, Nichols J, Ficz G, Krueger F, Oxley D, Santos F, Clarke J, Mansfield W, Reik W, Bertone P, et al. Resetting transcription factor control circuitry toward ground-state pluripotency in human. Cell 2014; 158:1254–1269.

[37] Putra VDL, Kilian KA, Knothe Tate ML. Biomechanical, biophysical and biochemical modulators of cytoskeletal remodelling and emergent stem cell lineage commitment. Communications Biology 2023 6:1 2023; 6:75-.

[38] Conway JRW, Isomursu A, Follain G, Härmä V, Jou-Ollé E, Pasquier N, Välimäki EPO, Rantala JK, Ivaska J. Defined extracellular matrix compositions support stiffness-insensitive cell spreading and adhesion signaling. Proc Natl Acad Sci U S A 2023; 120:e2304288120.

[39] Wang H, Luo X, Leighton J. Extracellular Matrix and Integrins in Embryonic Stem Cell Differentiation. Biochem Insights 2015; 8:15.

[40] Muncie JM, Weaver VM. The Physical and Biochemical Properties of the Extracellular Matrix Regulate Cell Fate. Curr Top Dev Biol 2018; 130:1.

[41] Sun Z, Guo SS, Fässler R. Integrin-mediated mechanotransduction. J Cell Biol 2016; 215:445.

[42] Wang X, Xiang Y, Yu Y, Wang R, Zhang Y, Xu Q, Sun H, Zhao ZA, Jiang X, Wang X, Lu X, Qin D, et al. Formative pluripotent stem cells show features of epiblast cells poised for gastrulation. Cell Research 2021 31:5 2021; 31:526–541.

[43] Weissbein U, Benvenisty N. rsPSCs: A new type of pluripotent stem cells. Cell Res 2015; 25:889–890.

[44] Morgani S, Nichols J, Hadjantonakis AK. The many faces of Pluripotency: in vitro adaptations of a continuum of in vivo states. BMC Dev Biol 2017; 17:7.

[45] Bi Y, Tu Z, Zhou J, Zhu X, Wang H, Gao S, Wang Y. Cell fate roadmap of human primed-to-naive transition reveals preimplantation cell lineage signatures. Nature Communications 2022 13:1 2022; 13:3147-.

[46] Zhi M, Zhang J, Tang Q, Yu D, Gao S, Gao D, Liu P, Guo J, Hai T, Gao J, Cao S, Zhao Z, et al. Generation and characterization of stable pig pregastrulation epiblast stem cell lines. Cell Research 2021 32:4 2021; 32:383–400.

